# California’s forest carbon offsets buffer pool is severely undercapitalized

**DOI:** 10.1101/2022.04.27.488938

**Authors:** Grayson Badgley, Freya Chay, Oriana S. Chegwidden, Joseph J. Hamman, Jeremy Freeman, Danny Cullenward

**Affiliations:** CarbonPlan, San Francisco, California, USA; Institute for Carbon Removal Law and Policy, American University, Washington, DC, USA

## Abstract

California operates a large forest carbon offsets program that credits carbon stored in forests across the continental United States and parts of coastal Alaska. These credits can be sold to buyers who wish to justify ongoing emissions, including in California’s cap-and-trade program. Although fossil CO_2_ emissions have effectively permanent atmospheric consequences, carbon stored in forests is inherently less durable because forests are subject to significant socioeconomic and physical risks that can cause temporarily stored carbon to be re-released into the atmosphere. To address these risks, California’s program is nominally designed to provide a 100-year guarantee on forest carbon claims based on a self-insurance program known as a buffer pool. Projects contribute credits to the buffer pool based on a suite of project-specific risk factors, with buffer pool credits retired as needed to cover carbon losses from events such as wildfire or drought. So long as the buffer pool remains solvent, the program’s permanence claim remains intact. Here, we perform an actuarial analysis of the performance of California’s buffer pool. We document how wildfires have depleted nearly one-fifth of the total buffer pool in less than a decade, equivalent to at least 95 percent of the program-wide contribution intended to manage all fire risks for 100 years. We also show that potential carbon losses from a single forest disease, sudden oak death, could fully encumber all credits set aside for disease and insect risks. These findings indicate that California’s buffer pool is severely undercapitalized and therefore unlikely to be able to guarantee the environmental integrity of California’s forest offsets program for 100 years.

## Introduction

Carbon offset programs have gained widespread adoption globally and have steadily increased in both size and scope. Over 2 billion credits were issued in the Clean Development and Joint Implementation carbon offset programs under the United Nations’ Kyoto Protocol, with more than one billion of these credits used in the European Union’s cap-and-trade program for greenhouse gasses — the largest single carbon market in the world (Ellerman et al., 2016).

Carbon offset quality concerns are central to climate policy accounting because offset credits increase the quantity of greenhouse gas emissions allowed within a legally binding policy system, in exchange for climate benefits claimed somewhere else (Cullenward & Victor, 2020; Erickson et al., 2014). We focus on California’s cap-and-trade program, which applies to about 75 percent of statewide emission sources. Polluters subject to the program must acquire and surrender pollution allowances issued under the cap-and-trade program. They can also use a limited number of carbon offsets that claim climate benefits in sectors outside of the carbon market.

After Europe decided to restrict the use of offsets in response to quality concerns, California’s cap-and-trade program emerged as one of the largest public markets for carbon offset credits (Haya et al., 2020). By the end of 2021, the California Air Resources Board had issued over 231 million offset credits, each worth 1 tCO_2_e (California Air Resources Board, 2022c), while regulated polluters in the linked cap-and-trade programs in California and Québec had surrendered just under 159 million offset credits to comply with program rules (Burtraw et al., 2022). Based on average fourth-quarter 2021 prices of about $16 per credit (California Air Resources Board, 2022d), California’s offset market capitalization was about $3.7 billion. Apart from its scale, California’s offsets program is relevant to study because forests across the United States are eligible to receive credits and because other jurisdictions, including the state of Washington, have proposed to adopt it in full in their own domestic climate policies.

The premise of carbon offsets is that they credit climate benefits that are equivalent to the emissions they justify (Carton et al., 2021; Gifford, 2020). This equivalency claim has been criticized on several dimensions, including: whether or not the offset projects credit non-additional, business-as-usual activities (Calel et al., 2021; Cames et al., 2016; Haya et al., 2020; Schneider, 2009); whether they cause emissions to shift or “leak” to other jurisdictions, rather than decrease net emissions on a global basis (Aukland et al., 2003; Schwartzman et al., 2021); and whether the baseline scenarios against which credits are issued represent realistic and credible counterfactuals (Badgley et al., 2022; Schneider, 2011; Schneider & Kollmuss, 2015; West et al., 2020).

Our study contributes to the carbon offsets literature by examining a separate issue known as permanence. We focus on California’s multi-billion dollar forest offsets program, which accounts for about 80 percent of total offset credits in the linked California-Québec cap-and-trade program (Burtraw et al., 2022). While California’s emissions limits only apply to polluters at the state level, forests throughout the continental United States and parts of coastal Alaska are eligible to receive offset credits, making the geographic footprint of the program much larger than California itself. Those forests, in turn, receive credits for implementing changes in forest management that promote carbon stocks in excess of regional common practice (California Air Resources Board, 2011, 2014, 2015). After verification, which includes on-the-ground field surveys, the state regulator issues offset credits that can then be used in California’s cap-and-trade program or sold on voluntary markets to justify CO_2_ emissions.

### Permanence

The permanence or durability of carbon stored in temporary carbon pools, such as the carbon stored in forests and soil, is an important dimension to consider when evaluating the efficacy of climate mitigation strategies (Kirschbaum, 2006; Matthews et al., 2022). Carbon dioxide emissions from fossil fuels have significant atmospheric impacts that last for hundreds to thousands of years (Archer et al., 2009; Joos et al., 2013) as well as effects that extend to geologic timescales (Pierrehumbert, 2014). In contrast, carbon stored in biological sinks is inherently more temporary and faces significant risks to permanence that are expected to increase in a changing climate (Anderegg et al., 2020).

The inherent impermanence of forest carbon introduces a fundamental asymmetry when used to offset effectively permanent fossil carbon emissions. The tension arises from the fiction that the physical climate benefits claimed by temporary carbon offsets are equivalent to the harms caused by ongoing pollution. In fact, the expected lifetime of biological carbon in temporary sinks like forests is necessarily shorter than the lifetime of fossil carbon in the atmosphere.

There is no easy way to resolve the tension between the distinct lifetimes of forest carbon and atmospheric CO_2_. California law requires that all carbon offsets be “permanent,” but does not define this term.^1^ The California Air Resources Board, which implements the state’s primary climate law, has interpreted “permanent” to require a minimum storage duration of 100 years.^2^ In turn, California’s forest offsets program explicitly accounts for the possibility that carbon temporarily stored by forests could be released back to the atmosphere — prior to the 100-year permanence period required by regulation — as a result of natural and non-natural risks (such as wildfire and bankruptcy, respectively). To achieve this goal, the California Air Resources Board developed a self-insurance mechanism called a buffer pool.

### Buffer pool

The purpose of California’s forest carbon buffer pool is to insure the permanence of the broader forest carbon offsets program. Whenever forest offset projects lose carbon due to factors that are outside of the landowner’s control — resulting in what are known as unintentional reversals — projects must conduct ground surveys to measure and report CO_2_ losses to the California Air Resources Board.^3^ Verified losses then trigger retirements from the buffer pool, such that one buffer pool credit is retired for every tCO_2_ lost, up to the total amount of project credits. Critically, credits in the buffer pool are cross-fungible: although contribution levels are based on a project’s individual risk factors, credits can be retired to account for any unintentional reversal. So long as the buffer pool remains solvent, the permanence claims of all credits in circulation remain intact.

California’s forest carbon buffer pool is capitalized by a share of offset credits issued by the California Air Resources Board. The number of credits contributed to the buffer pool is based on a series of project-specific risk factors. For example, in the currently applicable protocol, projects must contribute between 2 and 4 percent of their credits to account for wildfire risks, with lower levels allowed for projects that employ active wildfire management practices (California Air Resources Board, 2015). Projects must also contribute a fixed 3 percent of gross credits to account for disease- and insect-related mortality risks, along with another 3 percent for other catastrophic natural risks, such as wind, ice, and flood events. Finally, projects must also contribute between 1 and 9 percent of gross credits to account for various financial and management risks, such as the risk of bankruptcy, land use conversion, and excess timber harvesting.

We calculated contributions to the buffer pool through January 5, 2022 based on the program’s issuance table (California Air Resources Board, 2022a). To quantify projects’ variable contributions to the buffer pool to address wildfire risks, we used official project documentation to assemble the reported fire reversal factors for all 148 forest projects that had been issued credits prior to our study cutoff date, partially drawing on a previous data collection effort (Badgley et al., 2021). Because the disease and insect and other catastrophic natural risk components have fixed contribution rates, we inferred the non-natural component of the buffer pool as the difference between the total size of the buffer pool and the sum of wildfire, disease and insect, and other natural risk components. This approach allows us to calculate gross buffer pool contributions made prior to buffer pool retirements that have been verified in the past, thus avoiding double-counting of past reversals; alternative data sources only permit a current accounting net of past reversals (California Air Resources Board, 2022c).

Figure 1 reports buffer pool contributions through January 5, 2022. About 31.0 million credits were contributed in total, with about 56 percent from natural risks and 44 percent from non-natural risks. Contributions for natural risks from wildfire, disease and insects, and other catastrophic natural risks comprised about 19, 18, and 18 percent of the total, respectively. We use the composition of the buffer pool in Figure 1 as the basis for an actuarial analysis of the program’s performance in the face of forest carbon permanence risks to determine whether the buffer pool is adequately capitalized to insure against unintentional reversals over 100 years.

**Figure 1:**
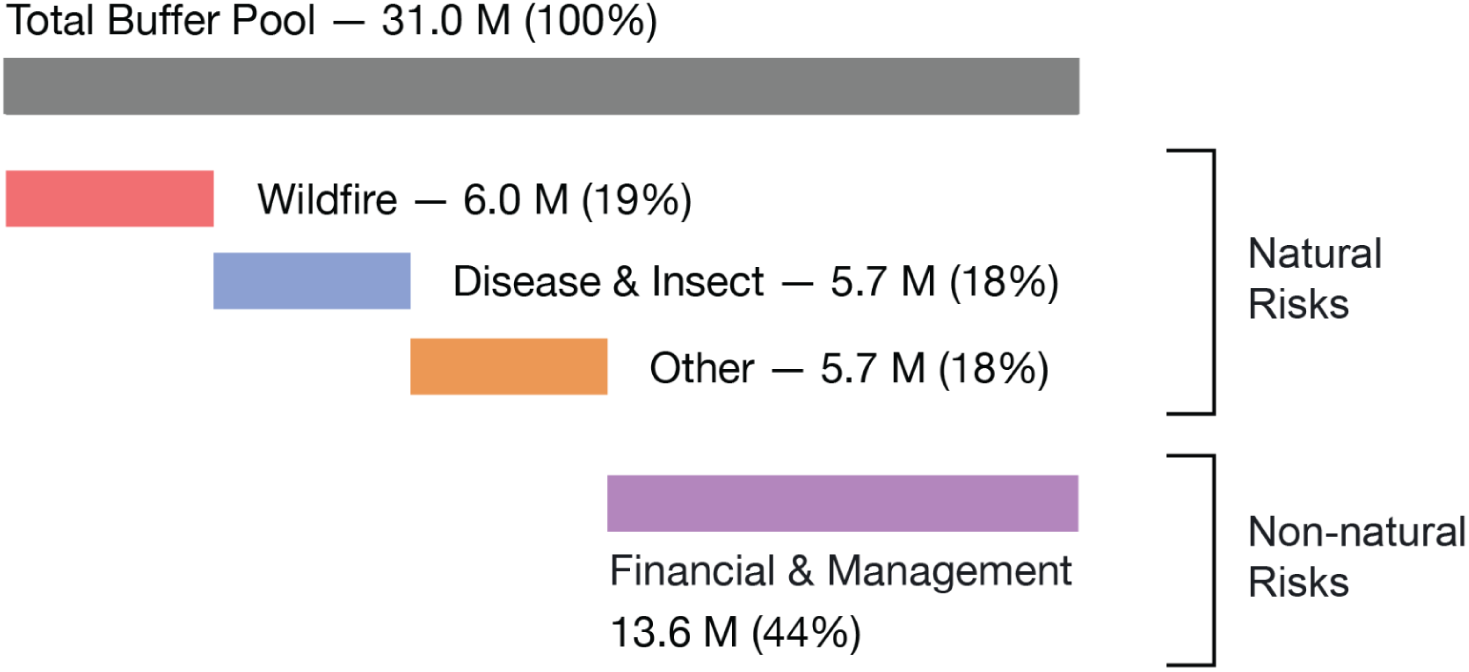
California forest buffer pool composition. Projects contribute a fraction of credited carbon to a communal buffer pool intended to insure against unintentional reversals. Under the 2015 forest carbon offsets protocol, projects contribute between 8.7 and 19.2 percent of credited carbon to the buffer pool. Each risk category is assigned a reversal risk rating. Wildfire risk (red) ranges between 2 and 4 percent of credited carbon, depending on if the project has an active management plan to mitigate wildfire risks.

Disease and insect risk (blue) is fixed at 3 percent. Other catastrophic risks (orange) like wind, ice, and flood events are also fixed at 3 percent. Financial and management risks (purple) vary by landowner type, such as private, public, or tribal lands, and conservation easement status. Percentage totals in the figure do not sum to 100 due to rounding.

Although these risk contributions are required by the protocol and, in practice, serve as the mechanism for ensuring the environmental integrity of California’s forest offsets program, we are unaware of any explicit analysis that justifies these numbers or the scientific basis of any of the buffer pool risk reversal ratings. Indeed, journalists interviewing the experts and policymakers who designed the buffer pool suggest that the risk ratings underpinning the buffer pool may have been the product of educated guesswork (Pontecorvo & Osaka, 2021).

We set out to evaluate how well this design appears to be working in light of recent unintentional carbon reversals in the program.

## Methods

We perform an actuarial analysis to evaluate the design and performance of California’s forest buffer pool mechanism. We focus on two specific risk categories covered by the buffer pool: (1) risks from wildfire and (2) risks from disease and insects. As described below, we estimate carbon losses directly from recent wildfire events and use scenario analysis to quantify potential future losses from a forest disease called sudden oak death. Together, these methods assess whether the buffer pool is large enough to provide 100 years of protection against these known risks.

Credits in the buffer pool are cross-fungible, meaning that any credit in the buffer pool can be retired to mitigate any unintentional reversal, no matter its cause. As a result, a risk-specific analysis can help identify whether a particular risk factor is undercapitalized or overcapitalized in the current buffer pool — that is, whether the number of credits set aside to protect against each risk factor is smaller or larger, respectively, than the expected loss from each risk factor. To the extent one or more components of the buffer is undercapitalized, the long-term solvency of the buffer pool as a whole depends on the remaining components being overcapitalized.

### Static portfolio analysis

We analyze a static portfolio of carbon offsets projects and a static view of the buffer pool, both of which are fixed in time as of the first week of January 2022 (California Air Resources Board, 2022a, 2022c). For the purposes of analysis, we hold constant the offset project portfolio as of this date and assume that the buffer pool is adequate to insure that fixed portfolio for the full crediting period plus 100 years.

Our analytical assumption that no new projects are added to the program past this point has two countervailing effects. First, we assume that no new contributions are made to the buffer pool from new projects. Second, we assume that no new liabilities are added to the program in the form of carbon credited in new projects. In reality, California’s forest offset program continues to add new projects, each of which contributes a share of the total credited carbon to the buffer pool (thereby increasing the size of the buffer pool). However, these new projects bring the possibility of future unintentional reversals (thereby increasing the liability exposure of the buffer pool). In the Discussion, we ask whether and how a dynamic portfolio analysis might affect the conclusions of our static analysis.

### Wildfires

Our wildfire analysis is designed to estimate the carbon losses associated with fires that have burned through offset projects to date, including the record-breaking 2020 and 2021 U.S. wildfire seasons. California’s offsets program gives projects affected by unintentional reversals up to 23 months to conduct field surveys and report total carbon losses.^4^ Thus, although unintentional reversals from the 2020 and 2021 wildfire seasons have already occurred, official estimates of the magnitude of these reversals are not yet available and the associated buffer pool credit retirements have not yet been made.

We estimate wildfire carbon losses by first identifying projects affected by wildfire and then quantifying the carbon reversals caused by those wildfires. We identify projects impacted by wildfire based on project perimeters downloaded from the California climate regulator (California Air Resources Board, 2022b). We then intersect the project perimeters with fire perimeters from the Monitoring Trends in Burn Severity (MTBS) database for fires through 2019 (MTBS Project, 2022) and the Wildland Fire Interagency Geospatial Services (WFIGS) Group for wildfires in 2020 and 2021 (National Interagency Fire Center, 2022). We identified six projects affected by wildfire, including two that have already reported verified reversals that have led to buffer pool credit retirements (CAR1046 and CAR1174) and four reversals that have not yet been verified by the regulator (ACR260, ACR273, CAR1102, and ACR255) (see Table 1).

**Table 1:** List of forest offset projects affected by wildfires.

| Project ID | Project name | Wildfire name | Year | Credits retired | Method |
| --- | --- | --- | --- | --- | --- |
| CAR1046 | Trinity Timberlands | Route Complex | 2015 | 847,895 | Observed |
| CAR1174 | Eddie Ranch | Ranch Fire | 2018 | 276,867 | Observed |
| ACR260 | Warm Springs | Lionshead | 2020 | Not yet reported | RAVG BA7 |
| ACR273 | Klamath East | Bootleg | 2021 | Not yet reported | RAVG BA7 |
| CAR1102 | Montesol | Glass;<br>Hennessey | 2020 | Not yet reported | Proxy |
| ACR255 | Colville | Chuweah Creek;<br>Summit Trail;<br>Whitmore | 2021 | Not yet reported | Proxy |

Once we have a list of offset projects and associated wildfires, we quantify expected carbon reversals based on the accounting rules used in the California forest offsets program. Verified carbon reversals for CAR1046 and CAR1174 are taken from official public reporting. Projected carbon reverals for the other four projects are based on a three-step process.

First, for each project we estimate on-site carbon stocks in the year prior to a wildfire impact. We begin with the most recently reported standing live trees (IFM-1) and standing dead trees (IFM-3). If there is a gap between project reporting dates and fire occurrences, we estimate changes to IFM-1 and IFM-3 using historical average growth rates from each project. In the case of ACR255, which has only filed two annual reports with CARB, we conservatively elect to base biomass losses on values of IFM-1 and IFM-3 reported in that project’s first crediting period, without accounting for forest growth that might contribute to higher fire losses. This step provides a conservative estimate of total living and dead carbon stocks at the time of fire, and thus the potential carbon losses from wildfire.

Second, we estimate wildfire mortality based on the U.S. Forest Service Rapid Assessment of Vegetation Condition 7-class percent change in basal area (RAVG BA7) data (Miller & Quayle, 2015; Miller & Thode, 2007). The RAVG BA7 data provide gridded estimates of the percent of basal area killed by fire, separated into seven classes of severity. Each severity class includes a minimum and maximum mortality estimate by percent basal area. For conservativeness, we assume the minimum mortality rates for the least severe classes (1 through 5) and vary the mortality rates for the most severe class (6 and 7) as discussed below. We use these data in two related methods to estimate changes in onsite carbon from wildfire for both standing live trees (IFM-1) and standing dead trees (IFM-3).

For two projects (ACR260 and ACR273), the U.S. Forest Service has published RAVG BA7 data. We intersect these mortality estimates with the project boundaries of each offset project and directly calculate area-weighted mortality and associated carbon losses under California’s accounting rules (see Figure 2). We call this the RAVG BA7 method.

**Figure 2:**
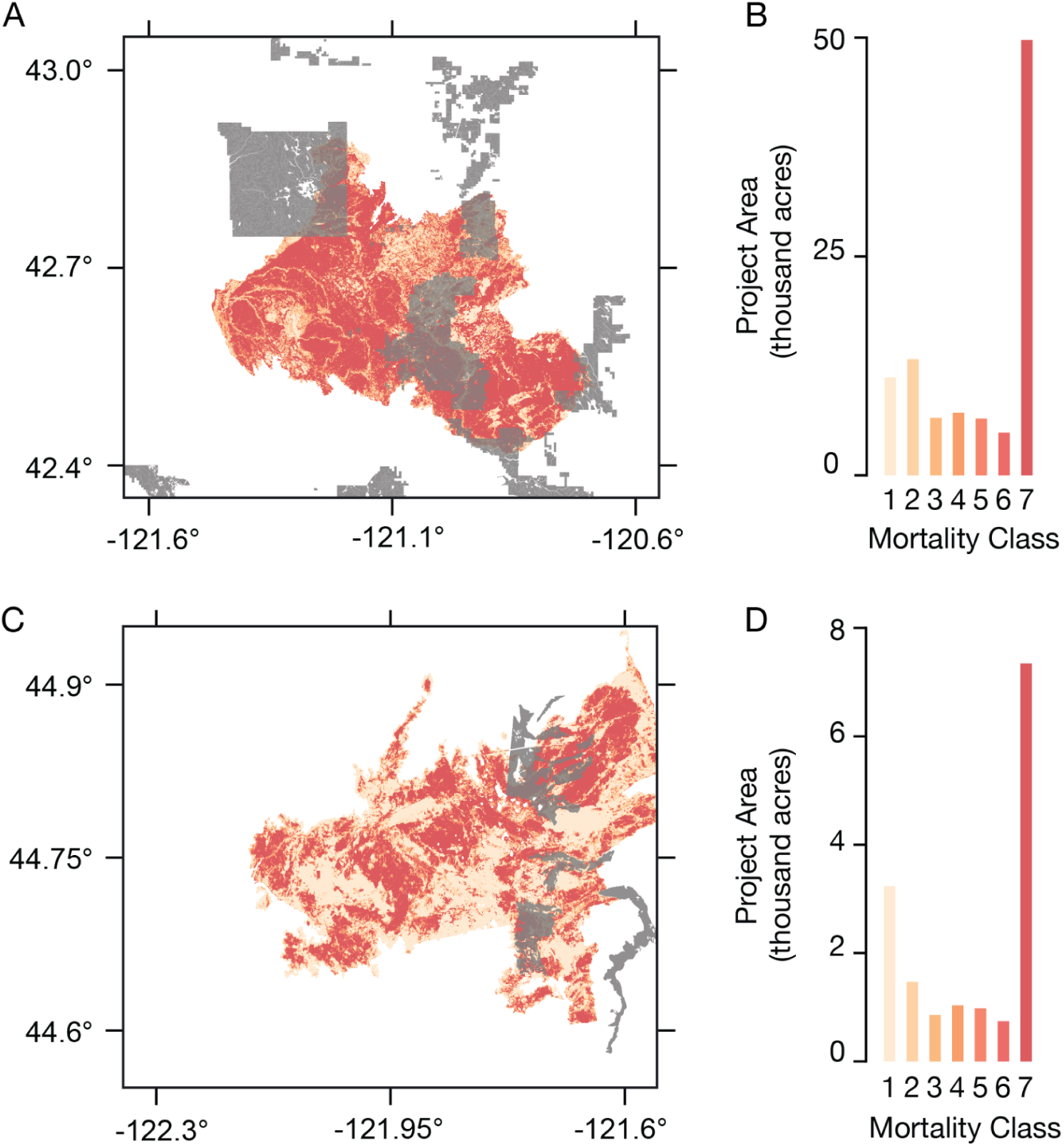
Estimated fire mortality in ACR260 and ACR273. Panel (A) shows the overlay between the Klamath East project (ACR273) in gray and the 2021 Bootleg fire, with a summary of the US Forest Service’s 7-class percent change in basal area data within the project area tabulated and shown in the same color shading in panel (B). Panels (C) and (D) show the same information for the Warm Springs project (ACR260) and the 2020 Lionshead fire.

For the other two projects (CAR1102 and ACR255), the U.S. Forest Service has not published RAVG BA7 data. We instead use proxy fires that have published RAVG BA7 data. In the case of CAR1102, we use proxy data from the Ranch fire, which burned CAR1174 in 2018 and resulted in a verified unintentional reversal that was covered by the buffer pool. This proxy is justified because CAR1102 and CAR1174 share similar forest types and climatic conditions, as reported in the projects’ official documentation. In the case of ACR255, we use proxy data from the 2015 North Star fire that burned through timberlands on the Colville Reservation that were originally eligible for inclusion in the ACR255 offsets project, but were burned while the project was still in development. Thus, the North Star fire burned through forests with a similar climate and species composition. Furthermore, we used climate anomaly data from the Oregon State University PRISM climate group to confirm that regional climatic conditions for eastern Washington were both drier and hotter in 2021, as compared to 2015, thus making the North Star fire a reasonable proxy for the 2021 fires that burned through ACR255. In both cases, we calculate the average mortality for the proxy fire (expressed as a percentage of basal area killed by wildfire) and apply that mortality factor to the observed burned area for each project during the 2020 and 2021 fire seasons. We call this the Proxy method.

Third, we estimate the amount of carbon transferred to wood products due to post-fire salvage harvest operations. California’s forest offsets program allows landowners to conduct post-fire salvage operations and deduct carbon stored in long-term wood products when calculating the size of unintentional reversals. This final step takes into account what fraction of fire-killed biomass is salvaged (which we call a “salvage fraction” in Table 2), the merchantable fraction of that salvaged biomass, and the fraction of merchantable biomass that is ultimately transferred to long-term wood products. The carbon stored in long-term wood products produced from salvaged biomass is then deducted from our estimates of biomass mortality, and thus reduces the magnitude of the unintentional reversal.

**Table 2:** Wildfire emission estimate methods.

| Scenario | Percent Mortality | Salvage Fraction | Standing dead (IFM-3) |
| --- | --- | --- | --- |
| Lower bound | Severity class 6: 75%<br>Severity class 7: 90% | 30% | No IFM-3 losses |
| Upper bound | Severity class 6: 90%<br>Severity class 7: 100% | 10% | IFM-3 losses included |

To report results across these six projects and their different conditions, we construct two representative scenarios to generate lower and upper bound estimates of wildfire reversals (Table 2).

For the lower bound scenario, we use minimum mortality rates as reported by the RAVG BA7 data (severity class 6: 75 percent of basal area killed, severity class 7: 90 percent of basal area killed); that post-fire salvage operations are extensive and therefore reported carbon reversals are lower (with a 30 percent salvage fraction); and we further exclude the loss of standing dead trees (IFM-3) despite carbon stored in standing dead trees being considered part of carbon reversals under the California program rules. Each of these assumptions produces a conservative bias to our estimates of wildfire reversals.

For the upper bound scenario, we assume maximum projected mortality rates in the RAVG BA7 data (severity class 6: 90 percent mortality, severity class 7: 100 percent mortality); that post-fire salvage operations are modest (with a 10 percent salvage fraction); and count the loss of both living and dead standing carbon as a reversal (consistent with the California program rules). Each of these assumptions is less conservative than in the lower bound scenario, though collectively they remain reasonable.

### Disease and insects

Our disease and insect analysis focuses on the California program’s exposure to unintentional reversals from a single pathogen and its anticipated effects on a single tree species that is prominent across California’s forest offsets project portfolio. Specifically, we analyze the expected impact of *Phytophthora ramorum*, an invasive pathogen that causes a forest disease called sudden oak death that disproportionately kills tanoak (*Notholithocarpus densiflorus*), a tree species endemic to the California and Oregon coast (Cobb et al., 2012, 2020).

In contrast to the wildfire analysis, which quantifies expected unintentional carbon reversals based on wildfires that have already occurred, our analysis here estimates the potential magnitude of future carbon losses. That is, our sudden oak death analysis considers possible losses that have not yet occurred, but which are liabilities that can be reasonably anticipated to encumber the buffer pool today.

The epidemiology of sudden oak death among tanoak is related to several environmental factors. Our analysis focuses on two of these factors. First, *P. ramorum* thrives and spreads most easily under cool, moist conditions (Cobb et al., 2012, 2020; Meentemeyer et al., 2011). Second, tanoak mortality risks are higher in the presence of California bay laurel (*Umbellularia californica*) because *P. ramorum* infects bay laurel trees but does not significantly harm them (Cobb et al., 2012; Kozanitas et al., 2022). Thus, the presence of bay laurel accelerates the spread of *P. ramorum* and significantly increases the odds of tanoak infection.

To assess potential future carbon losses from sudden oak death, we use an existing dataset of the species composition of individual California forest offset projects to identify projects that have credited carbon stored in tanoak trees (Badgley et al., 2021). For completeness, we updated these data to include five additional projects (CAR1313, CAR1329, CAR1330, CAR1339, CAR1368) that contain tanoak and were issued credits prior to our study cut-off date. For these 20 tanoak projects, we examined their filed paperwork to determine if California bay laurel was present, in any quantity, within their project boundaries. Furthermore, we conservatively exclude projects that initially enrolled in California’s forest offsets as Early Action projects. Our analysis also relies on estimates of the distribution of tanoak in California and Oregon provided by the LEMMA GNN dataset (Bell et al., 2018). We combine these data with gridded temperature data from PRISM to characterize population-level climatic conditions across the full geographic range of tanoak forests (PRISM Climate Group, 2016). Using these datasets and documented risk factors of tanoak mortality from sudden oak death, we develop three scenarios to explore potential future carbon losses.

Scenario A is a conservative scenario that explores possible tanoak mortality in relatively cooler climates. Using LEMMA, we identify the median mean annual temperature across all forests that contain tanoak throughout California and Oregon. We select only those forest offset projects that (1) contain tanoak and (2) have a mean annual average temperature, based on the centroid of project geometries and gridded PRISM data, that is less than the median of mean annual temperature across the full geographic range of tanoak. This method returns a list of 10 projects. We then assume that 50 percent of tanoak biomass will be lost to sudden oak death in these projects, and that no other tanoak mortality will occur elsewhere in the program.

Scenario B examines the impacts of sudden oak death, mediated by the presence of California bay laurel. Using official project documentation, we identify 17 projects in California’s forest offsets program that contain both tanoak and California bay laurel. We assume that 50 percent of tanoak biomass will be lost to sudden oak death in these projects, and that no other tanoak mortality will occur elsewhere in the program.

Scenario C attempts to characterize a more widespread mass mortality event. It assumes, for simplicity, that 80 percent of credited tanoak biomass will be lost to sudden oak death across the 20 projects in California’s program that contain this species. Although this scenario might appear aggressive, several ecologists warn of outcomes that are commensurate with this high degree of mortality (Cobb et al., 2012, 2020; Garbelotto, 2021).

## Results

Our analysis indicates that California’s forest offset buffer pool is severely undercapitalized. Estimated carbon losses from wildfires within the offset program’s first 10 years have depleted at least 95 percent of the contributions set aside to protect against all fire risks over 100 years. Even if we make the implausible assumption that no additional wildfires will impact forest offsets projects in California’s program, we nevertheless forecast that carbon reverals from historical fires will nearly drain and likely deplete the wildfire component of the buffer pool. Similarly, although the California program has not verified any carbon reversals associated with forest disease, the potential carbon losses associated with a single disease (sudden oak death) and its impacts on a single species (tanoak) is large enough to fully encumber the total credits set aside for all disease- and insect-related mortality over 100 years. Here again, even if we make the implausible assumption that no additional diseases or insects will cause forest carbon losses, sudden oak death alone has the potential to fully deplete the disease and insect component of the buffer pool.

### Wildfire

We identified six projects that have experienced significant wildfire events (Table 1). Two wildfire-induced reversals, which occurred in 2015 and 2018, have already been verified by CARB, resulting in the retirement of over 1.1 million offset credits from the buffer pool. After deducting these already retired credits, the net wildfire contribution to the buffer pool stands at 4.9 million credits. At least four additional projects have experienced significant wildfire events in subsequent years, with one project (ACR255) experiencing several large fires in 2021.

Using high-resolution, satellite-derived maps of wildfire-induced tree mortality generated by the U.S. Forest Service, we estimate committed carbon losses between 4.6 and 5.7 million credits resulting from the devastating 2020 and 2021 fire seasons. When combined with verified wildfire losses, we estimate wildfire has caused the reversal of between 5.7 and 6.8 million credits, representing between 95 percent and 114 percent of the credits earmarked to account for 100 years of portfolio-wide wildfire risks (Figure 3). In other words, we find that at least 95 percent of the wildfire component of the buffer pool that was intended to secure against the collective risk of fire reversals through the end of the 21st century has been depleted in less than a decade.

**Figure 3:**
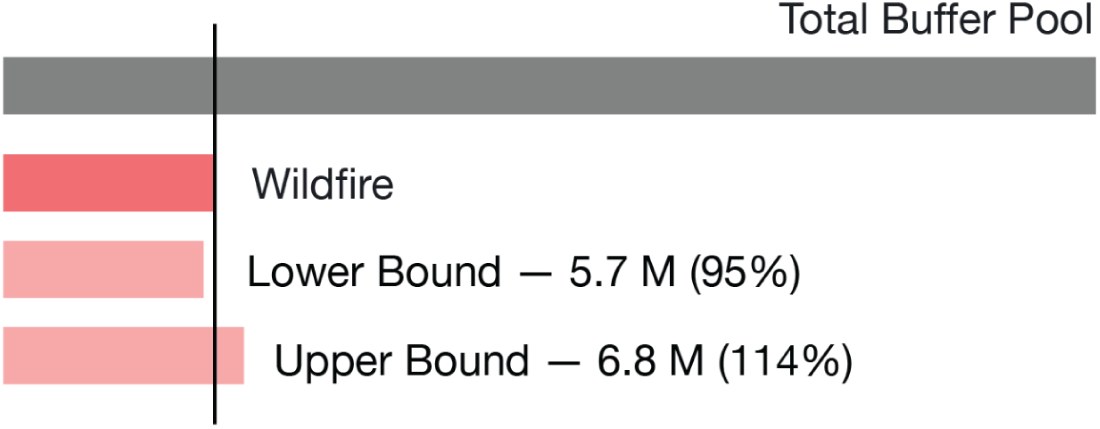
Carbon losses from wildfire. Estimated carbon losses from historical wildfires are large enough to deplete the entire buffer pool contribution set aside for all wildfire impacts over 100 years. We combine verified reversals reported by two projects with our estimates of carbon loss for four projects affected by wildfire. The Lower Bound scenario makes maximally conservative assumptions for wildfire mortality, post-fire salvage logging, and accounting treatment of standing dead carbon. The Upper Bound scenario relaxes the conservative assumptions for these variables.

**Figure 4:**
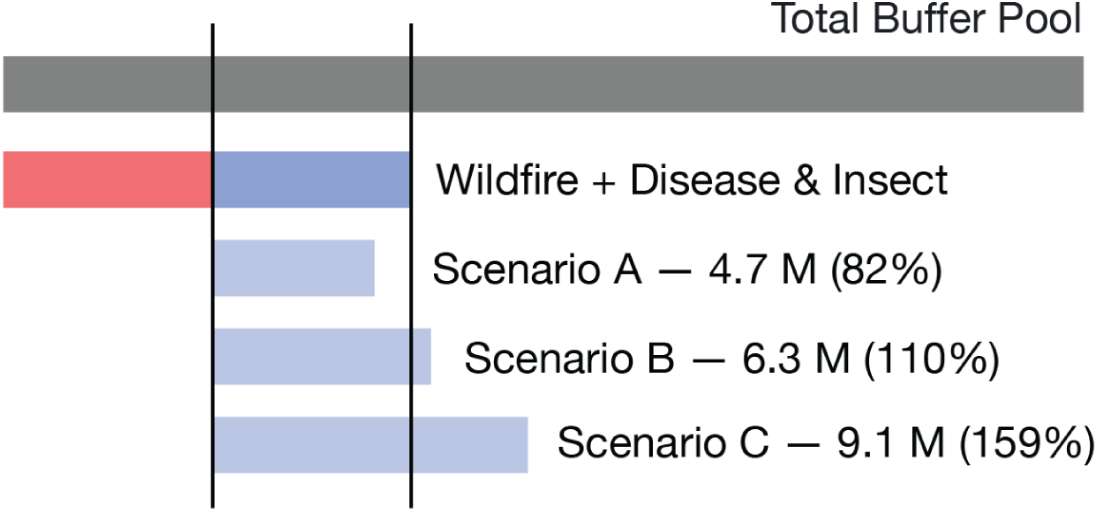
Carbon liabilities from sudden oak death. The potential liability from tanoak mortality caused by sudden oak death is large enough to encumber the entire buffer pool contribution set aside for all disease and insect risks over 100 years. Scenario A shows illustrative losses assuming 50 percent tanoak mortality, but only in relatively cool tanoak forests; Scenario B shows illustrative losses assuming 50 percent tanoak mortality, but only in forests that contain both tanoak and California bay laurel, a notable vector species of the pathogen *P. ramorum*; and Scenario C shows illustrative losses assuming 80 percent mortality of all credited tanoak in California’s forest carbon offsets program.

### Disease and insects

Our scenario analysis of potential tanoak mortality due to sudden oak death indicates that the disease and insect component of the buffer pool is likely undercapitalized. We identify 20 projects that collectively store nearly 14.2 million tCO_2_ in living tanoak stems, all of which are at risk to the pathogen *P. ramorum*. Our most conservative scenario (Scenario A) limits tanoak mortality to projects with relatively cool mean annual temperatures. It results in the expected loss of 4.7 million credits, or 82 percent of the disease and insect component of the buffer pool. Our moderate scenario (Scenario B) limits sudden oak death mortality to projects that also have a component of California bay laurel. It results in an expected loss of 6.3 million credits, or 110 percent of the disease and insect component of the buffer pool. Finally, the most aggressive scenario (Scenario C) assumes a loss of 80 percent of all tanoak biomass credited under California’s forest offsets program. It results in the expected loss of 9.1 million credits, or 159 percent of the disease and insect component of the buffer pool.

## Discussion

Assertions about the climate-equivalence of fossil CO_2_ emissions and forest carbon offsets depend on the permanence of the carbon stored in forests (Joppa et al., 2021). In turn, pricing these risks requires reasonable estimates of expected losses from disturbance. To be actuarially sound, a risk management approach needs to ensure that, on average, project-level risks match project-level insurance contributions.

Ensuring actuarial soundness is challenging enough when it comes to broad, regional estimates of risk factors. It may well be the case that reasonable estimates cannot be constructed to describe 100-year risks to forest carbon permanence in a changing climate (Williams & Jackson, 2007). Managing permanence risks is even more difficult because the composition of participating projects can be substantially different from regional averages due to selection effects (Badgley et al., 2022; Millard-Ball, 2013; Montero, 1999). For example, California assumes that, on average, no more than 3 percent of credited carbon will be lost to disease and insects; in the case of sudden oak death, however, the near-total loss of tanoak carbon is all but inevitable. The significant and preferential inclusion of tanoak projects in California’s program therefore acts as a net liability on the buffer pool. Accurately pricing permanence risks requires an evidence-based management approach that makes explicit assumptions about portfolio risks that are then evaluated against the actual composition of portfolio projects and updated over time with new information (Haya et al., 2020).

### Uncertainty and scenario analysis

Our wildfire analysis and disease and insect analysis require caveats with respect to how both analyses treat uncertainty and critical differences in the assumptions underlying each approach. We specifically developed our methods around a series of straightforward and conservative choices, with the overarching goal of facilitating expert interpretation of critical risks to the solvency of California’s forest offsets buffer pool.

The wildfire analysis represents an empirically grounded attempt to estimate carbon reversals from wildfires that have already occurred, but have not yet been validated by on-the-ground measurements pursuant to the offsets program’s rules. Because the analysis is underpinned by direct observations, we can attempt to account for uncertainty in the magnitude of estimated losses. RAVG BA7 data include a minimum and maximum mortality rate for each of seven basal area mortality classifications, but do not provide an explicit estimate of classification errors. Because the statistical ability to distinguish between severity classifications is greatest for the most severe classifications (e.g., 6 vs. 7), but relatively weaker for the least severe classifications (e.g., 1 vs. 2) (Miller & Quayle, 2015), we conservatively assumed that severity classifications 1 through 5 would cause the minimum mortality associated with their classification. We then explicitly varied assumed mortality for severity classification 6 (75 to 90 percent) and 7 (90 to 100 percent) to reflect uncertainty in mortality rates across severity classifications in which we had greater confidence. In addition, the two calculations we make for wildfire impacts based on proxy RAVG BA7 burn severity data depend on the reasonableness of the proxy fire we selected.

We elected to represent the limited explicit uncertainty contained in these methods, rather than ignore uncertainty altogether or introduce additional ad-hoc sources of uncertainty that lack any empirical basis. As a result, our lower and upper bound scenarios do not represent a statistically precise range of uncertainty, and instead are meant to communicate a plausible range of outcomes based on explicitly conservative methodological choices. For comparison, The Climate Trust (2021) estimates a total reversal of 6.8 million tCO_2_ across the Lionshead fire (ACR260), Bootleg fire (ACR273), and multiple fires that burned through ACR255 in 2021. We bound the losses from these same three events between 4.3 and 5.5 million tCO_2_, which indicates our approach — including the use of imperfect proxy methods — is conservative in relation to The Climate Trust’s calculations.

Our tanoak analysis is notably distinct from our wildfire estimates in that we are not estimating carbon losses that have already occurred, but rather projecting losses that might occur in the future. This evaluation is based entirely on constructing plausible future scenarios, which is inherently more speculative than an ex-post empirical calculation. Accordingly, we developed three scenarios based on simple, conservative assumptions. In the case of Scenario A, we modeled tanoak mortality only for projects that are in relatively cool climates and assume no sudden oak death mortality anywhere else. Similarly, in Scenario B we modeled tanoak mortality only in projects that also affirmatively report the presence of California bay laurel and assume no sudden oak death anywhere else. In practice, some projects may have even a handful of California bay laurel trees but not report them; and others will be located within a few hundred meters of California bay laurel trees that are outside of the project boundary. In both cases, our scenario construction excludes these likely sources of mortality. Finally, Scenario C represents a more catastrophic future that reflects concerns that sudden oak death could cause “devastating landscape-level mortality” (Cobb et al., 2020) and result in the complete extinction of tanoak populations at the local scale (Garbelotto, 2021).

### Buffer pool components

The undercapitalization of any one component of the buffer pool can in theory be compensated by overcapitalization in others. In our judgment, however, this outcome is implausible for California’s program on the basis of observable evidence. As discussed further below, the buffer pool design does not mention drought as a potential cause for forest death and carbon loss, with only a modest share of buffer pool credits (18 percent) set aside for all non-fire, non-insect, and non-disease carbon losses. The remaining component (44 percent) was designed to address financial and management risks associated with 100-year agreements that can be discharged in bankruptcy. We see no basis to believe that either the remaining natural risks or the financial and management risks are overcapitalized, and identify several reasons why these components are likely undercapitalized as well. Even if they were overcapitalized, the prospect of continuing wildfire impacts is potentially large enough to deplete the entire buffer pool in the decades to come (Herbert et al., 2020). We review carbon loss from wildfire, disease and insects, drought, and financial and management risks below.

### Wildfire

Historical carbon losses from wildfires are chiefly the result of the record-breaking 2020 and 2021 western U.S. fire seasons, which are unfortunately representative of the kinds of accelerating climate risks expected across the American West in a changing climate (Abatzoglou & Williams, 2016; Anderegg et al., 2022). Looking forward, wildfires are expected to grow in both size, intensity, and frequency as a result of anthropogenic climate change (Barbero et al., 2015). The increase in fire risk is dominated by rising global temperatures and, to date, there is little evidence that fuel-limitation will forestall the upward trend in annual burn area and increased fire intensity (Abatzoglou et al., 2021). Amidst this backdrop of increasing fire risk, it is critical to point out that California’s buffer pool makes no effort to account for the all-but-inevitable increase in fire risks as the Earth continues to warm. Nor does the program account for geographic variation in fire risks — forests in upstate New York are evaluated using the same risk reversal ratings that apply to forests in the arid American West (Anderegg et al., 2020; Pontecorvo & Osaka, 2021). Failure to acknowledge the increasing risk of wildfire means that California’s forest buffer pool is likely to experience mounting losses that far exceed its design criteria in the years and decades to come.

### Disease and insects

The continued spread of sudden oak death throughout California and southern Oregon indicates the need to seriously consider the potential effects of large-scale tanoak mortality events on the buffer pool. Using the sudden oak death (SOD) Blitz database (Garbelotto et al., 2014; Meentemeyer et al., 2015), a citizen science research project that tracks cases of *P. ramorum*, we calculated the distance between positive *P. ramorum* detections and forest offset projects. Of the 20 projects that report at least 5 percent tanoak basal area, the SOD Blitz database reports that one project (CAR1180) already has a positive *P. ramorum* sample within its boundaries. Another three projects are within 1 km of a positive detection (CAR1102, CAR1190, CAR1330); another two are within 5 km (CAR1313, CAR1339); another two are within 10 km (ACR182, ACR262); and all 20 of the tanoak projects we identified are within 50 km of a positive *P. ramorum* detection in the SOD Blitz database. The proximity of existing positive detections is a major concern for the long-term durability of carbon stored in tanoak biomass, as some researchers believe the *P. ramorum* is so widespread that it can no longer be eradicated and that active management is required to forestall widespread tanoak mortality (Cunniffe et al., 2016).

Critically, our analysis of the disease and insect component of the buffer pool only considers a single pathogen (*P. ramorum*) and the potential biomass losses of a single species (tanoak). There is growing evidence, however, that U.S. forests are likely to experience increased mortality pressure due to continuing globalization which brings with it more frequent opportunities for the introduction and spread of invasive pathogens. One recent analysis of Forest Inventory and Analysis data estimates that upwards of 40 percent of biomass stored in U.S. forests is subject to invasion by pathogens that are already established in North America, with the possibility of carbon loses to these pests and pathogens possibly exceeding 20.3 million tCO_2_ per year (Fei et al., 2019). In the same way that California’s buffer pool fails to account for changes in wildfire risks in the 21st century, it appears the buffer pool is likely unprepared to address these broader risks.

### Drought

The program’s buffer contribution for “other” natural risks is likely severely undercapitalized in the face of tree mortality risks from droughts that were not explicitly factored into its design. Current trends in the severity and frequency of drought-induced tree mortality have been characterized as both unexpected and truly without precedent (Hartmann et al., 2022).

Because these risks were not fully known when California designed its forest offsets program (Allen et al., 2010; Anderegg et al., 2013), it is likely the buffer pool is ill-suited to deal with this unique form of forest carbon disturbance. While uncertainties remain about the precise physiological causes of drought-induced mortality (Trugman et al., 2021), there is growing evidence that the effect of rising temperatures, superimposed on drought conditions, represents a novel threat to forests of all types, across all forested continents (Allen et al., 2015; Hammond et al., 2022). The connection between drought mortality and hotter climatic conditions is especially worrying given the current trajectory of global temperatures. From the standpoint of the forest carbon cycle, global-change-fueled droughts have the capacity to kill tens of millions of trees and profoundly alter the carbon balance of forested ecosystems (Sleeter et al., 2019; Stephens et al., 2018). Given that forest ecologists are still coming to terms with the unprecedented nature of global-change-fueled droughts as a unique disturbance agent in the 21st century, and had not fully grappled with the risks when California’s program was designed a decade ago, it is all but certain that the buffer pool does not adequately anticipate carbon losses due to drought.

### Financial and management risks

The buffer pool also includes a significant share of credits set aside to manage non-natural financial and management risks. One principal concern with these risks is that projects operate via long-term agreements that require landowners to comply with program rules and take actions to protect credited carbon. A handful of projects have elected to record these restrictions via easements that encumber their lands, and thus bind any future owners to those terms; however, the majority of forest offset projects operate via agreements between landowners and the California Air Resources Board. The liability for failing to perform can be discharged in bankruptcy, a risk that the California’s forest offsets protocol explicitly recognizes (California Air Resources Board, 2015, p. 133) and that was recently observed when a polluter in the cap-and-trade program successfully discharged its emissions liability in a bankruptcy proceeding.^5^ Thus, 100-year commitments made by individual or corporate landowners face significant risks. Together, the protocol sets aside a maximum of 9 percent of credits to cover nonperformance and bankruptcy risks for 100 years, although projects with conservation easements or those located on public or Tribal lands contribute only 1 percent.

These risk factors do not appear to be based on an explicit analysis of the probability of default or bankruptcy. It is worth noting, however, that California’s forest offsets protocol does not consider the creditworthiness of individual project proponents, which can be special-purpose limited liability companies. A probabilistic analysis of historical default risks based on credit ratings for companies tracked in the S&P 500 index suggests that AAA-rated firms have a 100-year default probability closer to 20 percent; BBB-rated firms, about 60 percent; and CCC-rated firms, nearly 100 percent (Mizdraković et al., 2015). These findings suggest that a contribution of 9 percent is unlikely to be adequate to protect against default risks over 100 years for all but the most financially sound entities.

### Static vs. dynamic portfolio analysis

Our results are based on a static portfolio analysis that freezes the carbon offsets projects enrolled in California’s program and the composition of its buffer pool at a fixed point in time, with the goal of asking whether the capitalization of the buffer pool is adequate to insure those assets (credits) in the face of their respective liabilities (observed or potential future carbon reversals). From this static point of view, our results indicate that the buffer pool is severely undercapitalized. Another approach would be to attempt to model a dynamic view of the buffer pool. Others have suggested that, if individual buffer pool components are undercapitalized, the addition of new projects with lower risks could help recapitalize the buffer pool by effectively cross-subsidizing older, riskier projects. For example, The Climate Trust (2021) argues that while the 2020 and 2021 wildfire seasons have led to significant reversals, the ongoing addition of new projects in relatively less fire-prone areas such as coastal Alaska could reduce the buffer pool’s undercapitalization. Although we recognize that the addition of new projects and new crediting periods will add new credits to the buffer pool, the extent to which every component of the buffer pool might be undercapitalized in its current design raises questions about the extent to which marginal additions of new projects — which ultimately include their own new liabilities — can mitigate an undercapitalized buffer pool. At a minimum, we suggest that claims about the ability of a dynamic portfolio to mitigate initial undercapitalization conditions should include a comprehensive analysis that looks at whether new projects are likely to outperform each risk factor, not just qualitative assertions that future projects might be less risky along one particular dimension.

### Credited vs. onsite carbon

The apparent disconnect between buffer pool capitalization and forest carbon permanence risks is amplified by an important but subtle asymmetry in the program’s carbon accounting rules. By design, California’s forest offset program awards a large initial tranche of credits to projects whose forests store more carbon than calculated regional averages (Badgley et al., 2022). Critically, most projects receive credit only for forest carbon storage in excess of projected 100-year average baseline levels, which must be equal to or higher than these regional averages (see Equation 5.1 in California Air Resources Board, 2015). In contrast, all standing trees contain carbon that is subject to reversal liability (see Equation 3.1 in California Air Resources Board, 2015). Thus, the potential carbon reversal liability insured by the buffer pool is substantially larger than the credited forest carbon that contributes to the buffer pool.

To give quantitative context for this dynamic, we examined a subset of 74 forest offset projects from a dataset assembled by Badgley et al. (2021). These projects collectively earned 121.2 million credits in their initial crediting periods, of which 20.3 million credits were contributed to the buffer pool. Although this implies a capitalization rate of 16.76 percent when compared against credited carbon, the onsite carbon stocks across these projects totaled 447.5 million tCO_2_. From the standpoint of onsite carbon liabilities, buffer pool contributions constitute only 4.54 percent of onsite carbon that is subject to reversal risks — less than a third of the capitalization implied by the nominal buffer pool contribution levels for these projects.

Asymmetry between the program’s calculation of assets and liabilities causes the buffer pool to be more highly leveraged and less resilient to reversals than it might at first appear. Seemingly modest reversals can trigger outsized buffer pool retirements, simply because the carbon liabilities (onsite carbon) strictly exceed carbon assets (credited carbon). This same asymmetry likely also affects buffer pools used by voluntary offset protocols, such as the Climate Action Reserve’s forest offsets protocol. Addressing the problem requires that permanence risks be properly calculated not on the basis of credited carbon, as is the case in California’s program, but on the basis of carbon subject to reversal liability — which we estimate is a factor of three larger. One solution would be to increase the share of credits projects must contribute to the buffer pool.

## Conclusion

We performed an actuarial analysis of California’s forest carbon offset buffer pool, a self-insurance program designed to ensure that credited forest carbon remains out of the atmosphere for at least 100 years. Whenever forest carbon offsets experience unintentional carbon reversals, such as losses from wildfire, credits are retired from the buffer pool to preserve the environmental integrity of all other forest carbon offset credits in the program. So long as the buffer pool is solvent, the program’s claim to keep credited forest carbon out of the atmosphere for at least 100 years remains intact.

From the program’s inception through our study cut-off date of January 5, 2022, a total of 31.0 million credits (13.4 percent) had been contributed to the buffer pool out of a total 231.5 million issued credits, such that the 31.0 million buffer pool credits insure a portfolio of 200.5 million credits against permanence risks. We reviewed how the buffer pool was constructed, but were unable to identify any explicit analysis that justifies the contribution levels required to address three categories of natural risks (wildfire, disease and insects, and other catastrophic risks) as well as a suite of financial and management risks.

To evaluate whether the buffer pool is adequately capitalized to address 100 years of risks to forest carbon permanence, we evaluated two specific permanence risks. First, we estimated total carbon losses from six historical wildfire events, based on verified reversals from two fires and our own estimates of four recent events. Second, we used scenario analysis to evaluate potential future carbon losses from sudden oak death. Our wildfire results show that at least 95 percent of the buffer pool contributions set aside to manage 100 years of wildfire risks had been depleted by the end of the 2021 wildfire season. Our sudden oak death scenarios indicate that carbon losses from this single disease could also fully encumber the credits set aside to protect against 100 years of risks from all disease- and insect-related mortality. These findings indicate that both the wildfire and the disease and insect buffer pool components are severely undercapitalized.

We explored whether there is reason to believe that any of the other components of the buffer pool are overcapitalized, such that the problems we identified with estimated actual losses from wildfire and potential future losses from sudden oak death might be mitigated by other, more conservative design choices. We did not find evidence that any buffer pool components are likely to be overcapitalized, and identified at least two mechanisms that suggest undercapitalization. First, the “other” catastrophic natural risks category likely underestimates the risk of drought-induced forest mortality, largely because the scope and scale of those risks are only now being fully realized. The evidence needed to accurately price drought risk simply was not available at the time when California’s forest offsets buffer pool was developed. Second, we showed how the total carbon stock exposed to reversal liability is about three times greater than the total carbon credited at projects enrolled in California’s program, illustrating how the buffer pool is significantly more leveraged as a result of broader exposure to permanence risks than is commonly understood.

Our analysis indicates that the wildfire and disease and insect components of the buffer pool are severely undercapitalized. None of the other buffer pool components shows any sign of being comparably overcapitalized, and indeed several well-documented factors suggest that forest mortality from wildfire, diseases and insects, and drought are likely to get significantly worse in the 21st century, not better. As a result, we conclude that California’s forest carbon buffer pool is severely undercapitalized and therefore unable to ensure that credited forest carbon remains out of the atmosphere for at least 100 years.

## Acknowledgements

We thank Matteo Garbelotto and Doug Schmidt for helpful discussions about tanoak and *P. ramorum*. Leander Anderegg offered feedback on the causes and consequences of drought-induced tree mortality. The USDA Forest Service RAVG Program helpfully provided a provisional copy of the Bootleg RAVG data, which assisted our initial analysis.

## Copyright and license

© 2022 The Authors. This document is licensed under a Creative Commons Attribution 4.0 International license, https://creativecommons.org/licenses/by/4.0/.

## Author contributions

Grayson and Danny designed the research. Grayson performed the research with input from all co-authors. Jeremy designed the figures. Danny and Grayson wrote the manuscript, with feedback from all co-authors.

## Conflict of Interest

The authors declare that this research was conducted in the absence of any commercial or financial relationships that could be construed as an actual or potential conflict of interest. Danny Cullenward is vice chair of California’s Independent Emissions Market Advisory Committee, but does not speak for the Committee here.

## Funding

This work received no specific source of funding. CarbonPlan’s funding is disclosed at https://carbonplan.org/funding.

## Open Source Software

We performed the core analysis using Python in a Pangeo cloud environment (Robinson et al., 2019). Our analysis relied on the following open software packages: GeoPandas (Jordahl et al., 2020); Jupyter (Kluyver et al., 2016); Matplotlib (Hunter, 2007); NumPy (Harris et al., 2020); Pandas (McKinney, 2010); rioxarray (Snow et al., 2022); Shapely (Gillies et al., 2007); tqdm (da Costa-Luis, 2019); and Xarray (Hoyer & Hamman, 2017). We also used rFIA (Stanke et al., 2020) and the R programming language (R Core Team, 2020) for a supplemental analysis not presented in the text.

## Data availability

The data underlying our analysis are available at https://doi.org/10.5281/zenodo.6465433 and an archive of the code is available at https://doi.org/10.5281/zenodo.6465461.

## Footnotes

1 California Health and Safety Code § 36562(d)(1) (added by Assembly Bill 32 in 2006).

2 California Code of Regulations, Title 17, § 95802 (defining “permanent” offsets as those that are either “irreversible” or have “mechanisms … to ensure that all credited reductions endure for at least 100 years”).

3 California Code of Regulations, Title 17, § 95983.

4 California Code of Regulations, Title 17, § 95983(b)(1).

5 California Air Resources Board v. La Paloma Generating Company, LLC, No. 1:17-CV-1698 (D. Del. Jul. 31, 2018).

## Notes

https://doi.org/10.5281/zenodo.6465433

https://doi.org/10.5281/zenodo.6465461

### Competing Interest Statement

The authors have declared no competing interest.

## References

Abatzoglou, J. T., Battisti, D. S., Williams, A. P., Hansen, W. D., Harvey, B. J., & Kolden, C. A. (2021). Projected increases in western US forest fire despite growing fuel constraints. Communications Earth & Environment, 2(1), 227. https://doi.org/10.1038/s43247-021-00299-0

Abatzoglou, J. T., & Williams, A. P. (2016). Impact of anthropogenic climate change on wildfire across western US forests. Proceedings of the National Academy of Sciences, 113(42), 11770–11775. https://doi.org/10.1073/pnas.1607171113

Allen, C. D., Breshears, D. D., & McDowell, N. G. (2015). On underestimation of global vulnerability to tree mortality and forest die-off from hotter drought in the Anthropocene. Ecosphere, 6(8), art129. https://doi.org/10.1890/ES15-00203.1

Allen, C. D., Macalady, A. K., Chenchouni, H., Bachelet, D., McDowell, N., Vennetier, M., Kitzberger, T., Rigling, A., Breshears, D. D., Hogg, E.H. (Ted), Gonzalez, P., Fensham, R., Zhang, Z., Castro, J., Demidova, N., Lim, J.-H., Allard, G., Running, S. W., Semerci, A., & Cobb, N. (2010). A global overview of drought and heat-induced tree mortality reveals emerging climate change risks for forests. Forest Ecology and Management, 259(4), 660–684. https://doi.org/10.1016/j.foreco.2009.09.001

Anderegg, W. R. L., Chegwidden, O., Badgley, G., Trugman, A. T., Cullenward, D., Abatzoglou, J., Hicke, J. A., Freeman, J., & Hamman, J. J. (2022). Future climate risks from stress, insects, and fire across US forests. Ecology Letters, in press.

Anderegg, W. R. L., Kane, J. M., & Anderegg, L. D. L. (2013). Consequences of widespread tree mortality triggered by drought and temperature stress. Nature Climate Change, 3(1), 30–36. https://doi.org/10.1038/nclimate1635

Anderegg, W. R. L., Trugman, A. T., Badgley, G., Anderson, C. M., Bartuska, A., Ciais, P., Cullenward, D., Field, C. B., Freeman, J., Goetz, S. J., Hicke, J. A., Huntzinger, D., Jackson, R. B., Nickerson, J., Pacala, S., & Randerson, J. T. (2020). Climate-driven risks to the climate mitigation potential of forests. Science, 368, eaaz7005. https://doi.org/10.1126/science.aaz7005

Archer, D., Eby, M., Brovkin, V., Ridgwell, A., Cao, L., Mikolajewicz, U., Caldeira, K., Matsumoto, K., Munhoven, G., Montenegro, A., & Tokos, K. (2009). Atmospheric Lifetime of Fossil Fuel Carbon Dioxide. Annual Review of Earth and Planetary Sciences, 37(1), 117–134. https://doi.org/10.1146/annurev.earth.031208.100206

Aukland, L., Costa, P. M., & Brown, S. (2003). A conceptual framework and its application for addressing leakage: The case of avoided deforestation. Climate Policy, 14.

Badgley, G., Freeman, J., Hamman, J. J., Haya, B., & Cullenward, D. (2021). California improved forest management offset project database (Version 1.0.0). https://doi.org/10.5281/zenodo.4630684

Badgley, G., Freeman, J., Hamman, J. J., Haya, B., Trugman, A. T., Anderegg, W. R. L., & Cullenward, D. (2022). Systematic over-crediting in California’s forest carbon offsets program. Global Change Biology, 28(4), 1433–1445. https://doi.org/10.1111/gcb.15943

Barbero, R., Abatzoglou, J. T., Larkin, N. K., Kolden, C. A., & Stocks, B. (2015). Climate change presents increased potential for very large fires in the contiguous United States. International Journal of Wildland Fire, 24(7), 892. https://doi.org/10.1071/WF15083

Bell, D. M., Gregory, M. J., Kane, V., Kane, J., Kennedy, R. E., Roberts, H. M., & Yang, Z. (2018). Multiscale divergence between Landsat- and lidar-based biomass mapping is related to regional variation in canopy cover and composition. Carbon Balance and Management, 13(1), 15. https://doi.org/10.1186/s13021-018-0104-6

Burtraw, D., Cullenward, D., Fowlie, M., Sutter, K. R., & Brown, R. (2022). 2021 Annual Report of the Independent Emissions Market Advisory Committee (p. 42). California Environmental Protection Agency. https://calepa.ca.gov/independent-emissions-market-advisory-committee/

Calel, R., Colmer, J., Dechezleprêtre, A., & Glachant, M. (2021). Do Carbon Offsets Offset Carbon? CESifo Working Paper No. 9368. https://doi.org/10.2139/ssrn.3950103

California Air Resources Board. (2011). Compliance Offset Protocol U.S. Forest Projects. https://ww2.arb.ca.gov/our-work/programs/compliance-offset-program/compliance-offset-protocols

California Air Resources Board. (2014). Compliance Offset Protocol U.S. Forest Projects. https://ww2.arb.ca.gov/our-work/programs/compliance-offset-program/compliance-offset-protocols

California Air Resources Board. (2015). Compliance Offset Protocol U.S. Forest Projects. https://ww2.arb.ca.gov/our-work/programs/compliance-offset-program/compliance-offset-protocols

California Air Resources Board. (2022a). ARB Offset Credit Issuance Table. https://ww2.arb.ca.gov/our-work/programs/compliance-offset-program/arb-offset-credit-issuance

California Air Resources Board. (2022b). ARBOC Issuance Map. https://webmaps.arb.ca.gov/ARBOCIssuanceMap/

California Air Resources Board. (2022c). Compliance Instrument Report—2021 Q4. https://ww2.arb.ca.gov/our-work/programs/cap-and-trade-program/program-data/compliance-instru ment-report

California Air Resources Board. (2022d). Summary of Transfers Registered in CITSS By California and Québec Entities During Fourth Quarter of 2021. https://ww2.arb.ca.gov/our-work/programs/cap-and-trade-program/program-data/summary-market-transfers-report

Cames, D. M., Harthan, D. R. O., Füssler, D. J., Lazarus, M., Lee, C. M., Erickson, P., & Spalding-Fecher, R. (2016). How additional is the Clean Development Mechanism? Öko-Institut e.V. https://ec.europa.eu/clima/sites/clima/files/ets/docs/clean_dev_mechanism_en.pdf

Carton, W., Lund, J. F., & Dooley, K. (2021). Undoing Equivalence: Rethinking Carbon Accounting for Just Carbon Removal. Frontiers in Climate, 3, 664130. https://doi.org/10.3389/fclim.2021.664130

Cobb, R. C., Filipe, J. A. N., Meentemeyer, R. K., Gilligan, C. A., & Rizzo, D. M. (2012). Ecosystem transformation by emerging infectious disease: Loss of large tanoak from California forests: Ecosystem transformation by disease. Journal of Ecology, 100(3), 712–722. https://doi.org/10.1111/j.1365-2745.2012.01960.x

Cobb, R. C., Haas, S. E., Kruskamp, N., Dillon, W. W., Swiecki, T. J., Rizzo, D. M., Frankel, S. J., & Meentemeyer, R. K. (2020). The Magnitude of Regional-Scale Tree Mortality Caused by the Invasive Pathogen Phytophthora ramorum. Earth’s Future, 8(7). https://doi.org/10.1029/2020EF001500

Cullenward, D., & Victor, D. G. (2020). Making Climate Policy Work. Polity.

Cunniffe, N. J., Cobb, R. C., Meentemeyer, R. K., Rizzo, D. M., & Gilligan, C. A. (2016). Modeling when, where, and how to manage a forest epidemic, motivated by sudden oak death in California. Proceedings of the National Academy of Sciences, 113(20), 5640–5645. https://doi.org/10.1073/pnas.1602153113

da Costa-Luis, C. O. (2019). tqdm: A Fast, Extensible Progress Meter for Python and CLI. Journal of Open Source Software, 4(37), 1277. https://doi.org/10.21105/joss.01277

Ellerman, A. D., Marcantonini, C., & Zaklan, A. (2016). The European Union Emissions Trading System: Ten Years and Counting. Review of Environmental Economics and Policy, 10(1), 89–107. https://doi.org/10.1093/reep/rev014

Erickson, P., Lazarus, M., & Spalding-Fecher, R. (2014). Net climate change mitigation of the Clean Development Mechanism. Energy Policy, 72, 146–154. https://doi.org/10.1016/j.enpol.2014.04.038

Fei, S., Morin, R. S., Oswalt, C. M., & Liebhold, A. M. (2019). Biomass losses resulting from insect and disease invasions in US forests. Proceedings of the National Academy of Sciences, 116(35), 17371–17376. https://doi.org/10.1073/pnas.1820601116

Garbelotto, M. (2021). Will emergent diseases decimate our forests? Feral Atlas. https://feralatlas.supdigital.org/poster/will-emergent-diseases-decimate-our-forests

Garbelotto, M., Maddison, E. R., & Schmidt, D. (2014). SODmap and SODmap Mobile: Two Tools to Monitor the Spread of Sudden Oak Death. Forest Phytophthoras, 4(1). https://doi.org/10.5399/osu/fp.4.1.3560

Gifford, L. (2020). “You can’t value what you can’t measure”: A critical look at forest carbon accounting. Climatic Change, 161(2), 291–306. https://doi.org/10.1007/s10584-020-02653-1

Gillies, S., Shenk, J., Taves, M., Van den Bossche, J., & Ostblom, J. (2007). Shapely: Manipulation and analysis of geometric objects. https://github.com/shapely/shapely

Hammond, W. M., Williams, A. P., Abatzoglou, J. T., Adams, H. D., Klein, T., López, R., Sáenz-Romero, C., Hartmann, H., Breshears, D. D., & Allen, C. D. (2022). Global field observations of tree die-off reveal hotter-drought fingerprint for Earth’s forests. Nature Communications, 13(1), 1761. https://doi.org/10.1038/s41467-022-29289-2

Harris, C. R., Millman, K. J., van der Walt, S. J., Gommers, R., Virtanen, P., Cournapeau, D., Wieser, E., Taylor, J., Berg, S., Smith, N. J., & others. (2020). Array programming with NumPy. Nature, 585(7825), 357–362.

Hartmann, H., Bastos, A., Das, A. J., Esquivel-Muelbert, A., Hammond, W. M., Martínez-Vilalta, J., McDowell, N. G., Powers, J. S., Pugh, T. A. M., Ruthrof, K. X., & Allen, C. D. (2022). Climate Change Risks to Global Forest Health: Emergence of Unexpected Events of Elevated Tree Mortality Worldwide. Annual Review of Plant Biology, 73(1), annurev-arplant-102820-012804. https://doi.org/10.1146/annurev-arplant-102820-012804

Haya, B., Cullenward, D., Strong, A. L., Grubert, E., Heilmayr, R., Sivas, D. A., & Wara, M. (2020). Managing uncertainty in carbon offsets: Insights from California’s standardized approach. Climate Policy, 20(9), 1112–1126. https://doi.org/10.1080/14693062.2020.1781035

Herbert, C., Stapp, J., Badgley, G., Anderegg, W. R. L., Cullenward, D., Hamman, J. J., & Freeman, J. (2020). Carbon offsets burning. CarbonPlan. https://carbonplan.org/research/offset-project-fire

Hoyer, S., & Hamman, J. (2017). xarray: N-D labeled arrays and datasets in Python. Journal of Open Research Software, 5(1). https://doi.org/10.5334/jors.148

Hunter, J. D. (2007). Matplotlib: A 2D Graphics Environment. Computing in Science & Engineering, 9(3), 90–95. https://doi.org/10.1109/MCSE.2007.55

Joos, F., Roth, R., Fuglestvedt, J. S., Peters, G. P., Enting, I. G., von Bloh, W., Brovkin, V., Burke, E. J., Eby, M., Edwards, N. R., Friedrich, T., Frölicher, T. L., Halloran, P. R., Holden, P. B., Jones, C., Kleinen, T., Mackenzie, F. T., Matsumoto, K., Meinshausen, M., … Weaver, A. J. (2013). Carbon dioxide and climate impulse response functions for the computation of greenhouse gas metrics: A multi-model analysis. Atmospheric Chemistry and Physics, 13(5), 2793–2825. https://doi.org/10.5194/acp-13-2793-2013

Joppa, L., Luers, A., Willmott, E., Friedmann, S. J., Hamburg, S. P., & Broze, R. (2021). Microsoft’s million-tonne CO2-removal purchase—Lessons for net zero. Nature, 597, 629–632. https://doi.org/10.1038/d41586-021-02606-3

Jordahl, K., Bossche, J. V. D., Fleischmann, M., Wasserman, J., McBride, J., Gerard, J., Tratner, J., Perry, M., Badaracco, A. G., Farmer, C., Hjelle, G. A., Snow, A. D., Cochran, M., Gillies, S., Culbertson, L., Bartos, M., Eubank, N., Maxalbert Bilogur, A., … Leblanc, F. (2020). geopandas/geopandas: V0.8.1 (v0.8.1) [Computer software]. Zenodo. https://doi.org/10.5281/ZENODO.3946761

Kirschbaum, M. U. F. (2006). Temporary Carbon Sequestration Cannot Prevent Climate Change. Mitigation and Adaptation Strategies for Global Change, 11(5–6), 1151–1164. https://doi.org/10.1007/s11027-006-9027-8

Kluyver, T., Ragan-Kelley, B., Pérez, F., Bussonnier, M., Frederic, J., Hamrick, J., Grout, J., Corlay, S., Ivanov, P., Abdalla, S., & Willing, C. (2016). Jupyter Notebooks—A publishing format for reproducible computational workflows. In F. Loizides & B. Schmidt (Eds.), Positioning and Power in Academic Publishing: Players, Agents and Agendas (pp. 87–90). IOS Publishing.

Kozanitas, M., Metz, M. R., Osmundson, T. W., Serrano, M. S., & Garbelotto, M. (2022). The Epidemiology of Sudden Oak Death Disease Caused by Phytophthora ramorum in a Mixed Bay Laurel-Oak Woodland Provides Important Clues for Disease Management. Pathogens, 11(2), 250. https://doi.org/10.3390/pathogens11020250

Matthews, H. D., Zickfeld, K., Dickau, M., MacIsaac, A. J., Mathesius, S., Nzotungicimpaye, C.-M., & Luers, A. (2022). Temporary nature-based carbon removal can lower peak warming in a well-below 2 °C scenario. Communications Earth & Environment, 3(1), 65. https://doi.org/10.1038/s43247-022-00391-z

McKinney, W. (2010). Data structures for statistical computing in python. Proceedings of the 9th Python in Science Conference, 445, 51–56.

Meentemeyer, R. K., Cunniffe, N. J., Cook, A. R., Filipe, J. A. N., Hunter, R. D., Rizzo, D. M., & Gilligan, C. A. (2011). Epidemiological modeling of invasion in heterogeneous landscapes: Spread of sudden oak death in California (1990–2030). Ecosphere, 2(2), art17. https://doi.org/10.1890/ES10-00192.1

Meentemeyer, R. K., Dorning, M. A., Vogler, J. B., Schmidt, D., & Garbelotto, M. (2015). Citizen science helps predict risk of emerging infectious disease. Frontiers in Ecology and the Environment, 13(4), 189–194. https://doi.org/10.1890/140299

Millard-Ball, A. (2013). The trouble with voluntary emissions trading: Uncertainty and adverse selection in sectoral crediting programs. Journal of Environmental Economics and Management, 65(1), 40–55. https://doi.org/10.1016/j.jeem.2012.05.007

Miller, J. D., & Quayle, B. (2015). Calibration and Validation of Immediate Post-Fire Satellite-Derived Data to Three Severity Metrics. Fire Ecology, 11(2), 12–30. https://doi.org/10.4996/fireecology.1102012

Miller, J. D., & Thode, A. E. (2007). Quantifying burn severity in a heterogeneous landscape with a relative version of the delta Normalized Burn Ratio (dNBR). Remote Sensing of Environment, 109(1), 66–80. https://doi.org/10.1016/j.rse.2006.12.006

Mizdraković, V., Stanišić, N., Popovčić-Avrić, S., & Ðenić, M. (2015). Predicting the Lifespan of a Company: An Important Factor for Capital Reallocat. Proceedings of the International Scientific Conference - Synthesis 2015, 394–399. https://doi.org/10.15308/Synthesis-2015-394-399

Montero, J. (1999). Voluntary Compliance with Market-Based Environmental Policy: Evidence from the U. S. Acid Rain Program. Journal of Political Economy, 107(5), 998–1033. https://doi.org/10.1086/250088

MTBS Project. (2022). MTBS Data Access. https://mtbs.gov/direct-download

National Interagency Fire Center. (2022). WFGIS - Wildland Fire Permiters Full History. https://data-nifc.opendata.arcgis.com/datasets/

Pierrehumbert, R. T. (2014). Short-Lived Climate Pollution. Annual Review of Earth and Planetary Sciences, 42(1), 341–379. https://doi.org/10.1146/annurev-earth-060313-054843

Pontecorvo, E., & Osaka, S. (2021, October 27). California is banking on forests to reduce emissions. What happens when they go up in smoke? Grist. https://grist.org/wildfires/california-forests-carbon-offsets-reduce-emissions/

PRISM Climate Group. (2016). PRISM 30-year Climate Normals. https://prism.oregonstate.edu/normals/

R Core Team. (2020). R: A language and environment for statistical computing [Manual]. https://www.R-project.org/

Robinson, N. H., Hamman, J., & Abernathey, R. (2019). Seven Principles for Effective Scientific Big-DataSystems. https://doi.org/10.48550/ARXIV.1908.03356

Schneider, L. (2009). Assessing the additionality of CDM projects: Practical experiences and lessons learned. Climate Policy, 9(3), 242–254. https://doi.org/10.3763/cpol.2008.0533

Schneider, L. (2011). Perverse incentives under the CDM: An evaluation of HFC-23 destruction projects. Climate Policy, 11(2), 851–864. https://doi.org/10.3763/cpol.2010.0096

Schneider, L., & Kollmuss, A. (2015). Perverse effects of carbon markets on HFC-23 and SF6 abatement projects in Russia. Nature Climate Change, 5(12), 1061–1063. https://doi.org/10.1038/nclimate2772

Schwartzman, S., Lubowski, R. N., Pacala, S. W., Keohane, N. O., Kerr, S., Oppenheimer, M., & Hamburg, S. P. (2021). Environmental integrity of emissions reductions depends on scale and systemic changes, not sector of origin. Environmental Research Letters, 16(9), 091001. https://doi.org/10.1088/1748-9326/ac18e8

Sleeter, B. M., Marvin, D. C., Cameron, D. R., Selmants, P. C., Westerling, A. L., Kreitler, J., Daniel, C. J., Liu, J., & Wilson, T. S. (2019). Effects of 21st-century climate, land use, and disturbances on ecosystem carbon balance in California. Global Change Biology, 25(10), 3334–3353. https://doi.org/10.1111/gcb.14677

Snow, A. D., Brochart, D., Raspaud, M., Bell, R., Chegini, T., Amici, A., Braun, R., Annex, A., Hoese, D., Bunt, F., GBallesteros Hamman, J., Zehner, M., Cordeiro, M., RichardScottOZ Henderson, S., Miller, S., Badger, T. G., Augspurger, T., & Pmallas. (2022). corteva/rioxarray: 0.10.3 Release (0.10.3) [Computer software]. Zenodo. https://doi.org/10.5281/ZENODO.6399263

Stanke, H., Finley, A. O., Weed, A. S., Walters, B. F., & Domke, G. M. (2020). rFIA: An R package for estimation of forest attributes with the US Forest Inventory and Analysis database. Environmental Modelling & Software, 127, 104664. https://doi.org/10.1016/j.envsoft.2020.104664

Stephens, S. L., Collins, B. M., Fettig, C. J., Finney, M. A., Hoffman, C. M., Knapp, E. E., North, M. P., Safford, H., & Wayman, R. B. (2018). Drought, Tree Mortality, and Wildfire in Forests Adapted to Frequent Fire. BioScience, 68(2), 77–88. https://doi.org/10.1093/biosci/bix146

The Climate Trust. (2021). California ARB buffer mitigates current wildfire risk to forest carbon projects. https://climatetrust.org/california-arb-buffer-mitigates-current-wildfire-risk-to-forest-carbon-projects/

Trugman, A. T., Anderegg, L. D. L., Anderegg, W. R. L., Das, A. J., & Stephenson, N. L. (2021). Why is Tree Drought Mortality so Hard to Predict? Trends in Ecology & Evolution, 36(6), 520–532. https://doi.org/10.1016/j.tree.2021.02.001

West, T. A. P., Börner, J., Sills, E. O., & Kontoleon, A. (2020). Overstated carbon emission reductions from voluntary REDD+ projects in the Brazilian Amazon. Proceedings of the National Academy of Sciences, 117(39), 24188–24194. https://doi.org/10.1073/pnas.2004334117

Williams, J. W., & Jackson, S. T. (2007). Novel climates, no-analog communities, and ecological surprises. Frontiers in Ecology and the Environment, 5(9), 475–482. https://doi.org/10.1890/070037

